# Hippocampal Pattern Separation Supports Reinforcement Learning

**DOI:** 10.1101/293332

**Authors:** Ian Ballard, Anthony D. Wagner, Samuel M. McClure

## Abstract

Animals rely on learned associations to make decisions. Associations can be based on relationships between object features (e.g., the three-leaflets of poison ivy leaves) and outcomes (e.g., rash). More often, outcomes are linked to multidimensional states (e.g., poison ivy is green in summer but red in spring). Feature-based reinforcement learning fails when the values of individual features depend on the other features present. One solution is to assign value to multifeatural conjunctive representations. We tested if the hippocampus formed separable conjunctive representations that enabled learning of response contingencies for stimuli of the form: AB+, B-, AC-, C+. Pattern analyses on functional MRI data showed the hippocampus formed conjunctive representations that were dissociable from feature components and that these representations influenced striatal PEs. Our results establish a novel role for hippocampal pattern separation and conjunctive representation in reinforcement learning.

---

Most North American hikers develop a reflexive aversion to poison ivy, which causes a painful rash, and learn to recognize its compound leaf with three leaflets that is green in summer and red in spring and autumn. The relationship between color and season distinguishes poison ivy from other plants like boxelder, which looks similar but is green in spring. Such learning problems are challenging because similar conjunctions of features can require different responses or elicit different predictions about future events. Responses and predictions also depend on the status of other features or context. In such problems, simple feature-response learning is insufficient and representations that include multiple features (leaf shape, color, season) must be learned.

Learning in the brain encode qualitatively distinct representations depending on the brain systems being considered. Theories posit that reinforcement learning likewise operates over multiple types of representations^1^. Theoretical and empirical work suggest the hippocampus rapidly forms conjunctive representations of arbitrary sets of co-occurring features^2^, making the hippocampus critical for episodic memory^3^. During encoding of conjunctive representations, hippocampal computations establish minimal representational overlap between traces of events with partially shared features, a process called pattern separation^4,5^, which reduces interference between experiences with overlapping features. One solution to multifeatural learning problems that require stimuli with overlapping features to be associated with different outcomes is to encode neurally separable conjunctive representations, putatively through hippocampal-dependent computations, and to assign value to each “separated” representation, putatively through hippocampal-striatal interactions. The same circuit and computational properties that make the hippocampus vital for episodic memory can also benefit striatal-dependent reinforcement learning by providing separated conjunctive representations over which value learning can occur.

Stimulus-response learning occurs by the incremental adjustment of synapses on striatal neurons^6^. Thalamic and sensory cortical inputs encode single stimuli, such as a reward-associated light, and are strengthened in response to phasic dopamine reward prediction errors (PEs)^7–9^. This system allows for incremental learning about individual feature values. Although the hippocampus is not critical for associating value with individual features or items^10^, it provides dense input to the striatum^11^. These synapses are strengthened by phasic dopamine release via D1 receptors^12^ and might represent conjunctions of features distributed in space or time^6^. We used a non-spatial, probabilistic stimulus-response learning task including stimuli with overlapping features to test the role of the hippocampus and its interaction with the striatum in value learning over conjunctive codes. We hypothesized that hippocampal pattern separation computations and hippocampal-to-striatal projections would form a conjunctive-value learning system that worked in tandem with a feature-value learning system implemented in sensory cortical-to-striatal projections.

We compared hippocampal representational codes to those of four other cortical areas that could contribute to learning in our task: perirhinal (PRc) and parahippocampal (PHc) cortices, inferior frontal sulcus (IFS), and medial orbitofrontal cortex (mOFC). The PRc and PHc gradually learn representations of individual items^13,14^. Cortical learning is generally too slow to form representations linking multiple items^2^, and pattern separation likely depends on hippocampal computations^5^. We therefore predicted PRc and PHc would not form pattern-separated representations of conjunctions with overlapping features. The IFS supports the representation of abstract rules^15,16^ that often describe conjunctive relationships (e.g., “respond to stimuli with both features A and B” ^17^, but our task included design features intended to biased subjects away from rule-based learning, and thus we predicted the IFS would not form pattern-separated representations of conjunctions. Finally, the mOFC is involved in outcome evaluation^18–20^, has been proposed to provide a state representation in learning tasks^21^ and receives dense medial temporal lobe inputs^22^. Due to its prominent role in reward processing, we predicted that the mOFC representations would be organized around the probability of reward associated with the stimuli, rather than idiosyncratic stimulus features. We designed our task and analyses to test for a hippocampal role in encoding conjunctive representations that serve as inputs for striatal associative learning.

## 2 RESULTS

### 2.1 Behavioral Results

Subjects learned stimulus-outcome relationships that required the formation of conjunctive representations. Our task was based on the “simultaneous feature discrimination” rodent behavioral paradigm^23^. Task stimuli consisted of four feature configurations (AB, AC, B, C). We used a speeded reaction time (RT) task in which a target “go” stimulus was differentially predicted by the four stimuli (Figure 1). AB and C predicted the target 70% of the time and B and AC 30% of the time. To earn money, subjects pressed a button within a limited response window after target onset. The response window was set adaptively for each subject and adjusted over the course of the run. Each feature was associated with the target 50% of the time, but stimuli were more (70%) or less (30%) predictive of the target. Optimal performance required learning the value of stimuli as distinct conjunctions of features (i.e., *conjunctive representations*).

**Figure 1.**
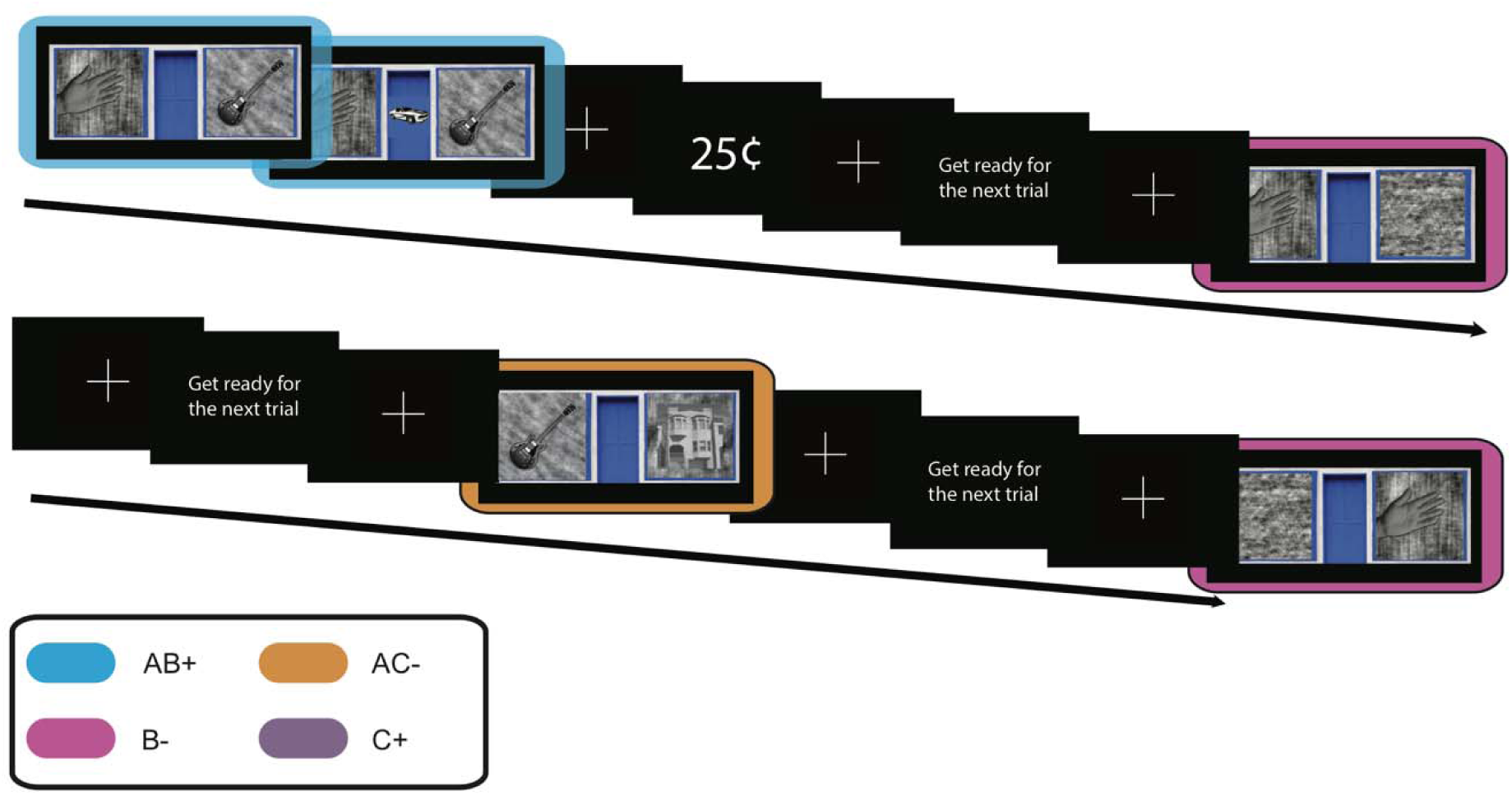
Task design. AB+, B- and AC-trials. The target appeared at fixation 600 ms after stimulus onset. Stimuli were always presented for 2000 ms. Feedback indicated whether subjects responded quickly enough to earn a reward. Note that faces were used instead of objects, but we have replaced the image of a face with an image of a guitar to conform to Biorxiv policy.

We first tested whether subjects learned predictive relationships between the stimuli and the target. Subjects were faster in responding to the target when it followed stimuli that were more reliably linked to target onset (AB+ and C+) than those that were less reliably linked (AC- and B-), *F*(1,26) = 13, *p* = .001, = .08, Figure 2b. Further, reaction times for target-predictive stimuli decreased over the course of each run, *Z* = −2.04, *p* = .041, mixed effects model with subject as a random intercept, and the adaptive RT threshold decreased as well (Figure S1). As a result of faster RTs, subjects had a higher hit rate for target stimuli, *F*(1,26) = 43, *p* < .001, 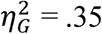, Figure 2a. In addition, subjects were more likely to make false alarms to target-predictive stimuli when the target did not appear, *F*(1,26) = 34, *p* = .001, 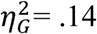, Figure 2c.

**Figure 2.**
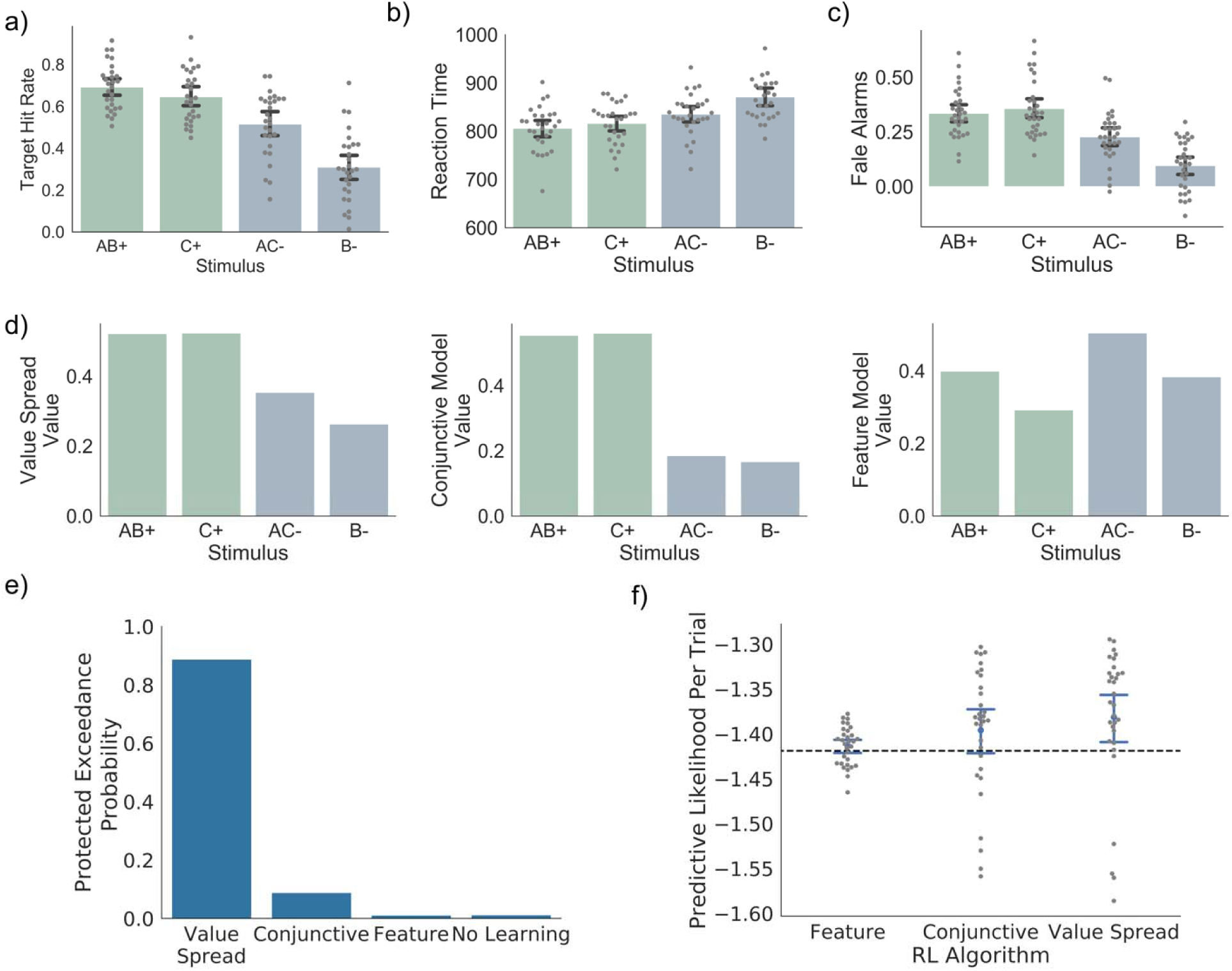
Behavior and Modeling Results. a.) Proportion of target trials in which the subject responded quickly enough to the target to earn a reward. Stimuli that are associated with the target (AB+, C+; green) have a higher hit rate than those that are not (AC-, B-; blue). In addition, stimuli with single features (B, C) are associated with a lower hit rate than those with two features (AB, AC). Further, this feature effect interacts with the target outcome effect. b.) Reaction times for each of the stimulus types. Reaction times are faster for stimuli associated with a target. The use of an adaptive RT threshold made relatively small differences in reaction times between conditions result in larger differences in hit rate in A. c.) False alarms for each stimulus type. Subjects are more likely to respond when no target occurred for stimuli that are associated with the target. d.) Value estimates from the Value Spread, Conjunctive and Feature models. The Conjunctive model shows an effect of target, such that AB+ and C+ have higher value than B- and AC-. The Feature model shows an effect of features, such that AB+ and AC-have a higher value than B+ and C-. Only the Value Spread model shows both the effect of target and the interaction between target and the number of features (present in A, B and C). e) Bayesian random effects model comparison showed the Value Spread RL model most likely accounted for behavior. Protected exceedance probabilities sum to 1 across models and because they express a group random-effects measure, there are no error bars. f)Cross-validation model comparison showed the Value Spread RL model best predicted unseen data. Log predictive likelihoods closer to 0 indicate better performance. Likelihoods are expressed per trial to normalize across differences in the number of responses between subjects. The dashed black line indicates the performance of the null model.

We aimed to identify the mechanism by which subjects learned stimulus-outcome relationships by fitting four computational models:

1) No Learning Model: subjects ignored predictive information and responded as fast as possible after target.

2) Feature RL: subjects learned values for individual features but not conjunctions. For multifeatural cues, value was updated for each feature.

3) Conjunctive RL: subjects learned values for each distinct stimulus. Value was updated for one representation on each trial (for “AB”, value updated for AB but not A or B).

4) Value Spread RL: subjects learned values of stimuli but confused stimuli that shared common features (e.g., AB and B). This model spreads value updates between stimuli that shared features (for AB trial, some of value update was applied to B).

Stimuli that are highly predictive of targets are associated with faster responses, permitting us to fit each model to the RT data. We first compared the Conjunctive model, which implements the experimenter-defined optimal task strategy, with the Feature and No Learning models. The Feature model uses a simpler and commonly used learning strategy, also referred to as “function approximation” or “feature weight” learning^24^. Although feature learning is not adaptive for the task, it may be the default learning strategy and exert an influence on learning^25^. Both the Feature and Conjunctive models have 2 free parameters: learning rate (*α*), and a regression weight relating values to reaction times (*β*). We assessed model fits using a cross-validated predictive likelihood method. The Conjunctive model outperformed the No Learning model, *T* = 136, *p* = .028, Wilcoxon test, Figure 1c, but was only marginally better than the Feature Model, *T* = 158, *p* = .08, Figure 1c. We next assessed the relative fits of these three models with a random-effects Bayesian procedure that gives probabilities that each model would generate the data of a random subject^26^. We found the most likely model was the Conjunctive model (protected exceedance probabilities (pEP): Conjunctive 92.3%, Feature 3.9%, No Learning 3.7%), which suggests subjects learned predictive relationships. Overall, there was mixed evidence in support of learning about conjunctions.

We reasoned that these results could be explained by subjects forming conjunctive representations and simultaneously learning the predictive value of individual features. This behavior could arise if hippocampal pattern separation was partially effective in encoding distinct representations for each stimulus^5^ and/or stimulus representations in the hippocampus and feature representations in cortex were simultaneously reinforced during learning^27^. We fit a Value Spread model that allowed for value updates to spread between stimuli with overlapping features. A parameter ω specifies the degree to which value updates spread to other stimuli with shared features, resulting in 3 free parameters (ω, *α, β*). This model outperformed both the Conjunctive, *T* = 124, *p* = .015, and Feature models, *T* = 115, *p* = .009; Wilcoxon tests, Figure 2f, on the cross-validation analysis. Bayesian model comparison confirmed the Value Spread model was the most likely model, pEP: 89.9%, Figure 2e. Values from the Value Spread model were anticorrelated with reaction times, mean *r* = .43, *t*(30) = 20, *p* < .001, Fisher corrected. In addition, fitted regression weights for the Value Spread model were significantly less than 0, *T* = 64, *p* < .001, Wilcoxon test, indicating that stimuli more strongly associated with the target are associated with faster reaction times. Fits of spread parameter ω (mean: .44, SD: .25, Table S1) indicated that for any given value update to the current stimulus (e.g., AB), about half that update was also applied to overlapping stimuli (e.g., B). Note mixed conjunctive and feature learning could also arise if the predictions of independent feature and conjunctive learnings systems were mixed at the time of outcome prediction. However, we were unable to reliability to fit such a model to our data. Therefore, we cannot adjudicate between spreading of value updates during learning versus mixed predictions of independent systems. Nonetheless, both models share the core feature that subjects form conjunctive representations of multifeatural stimuli but also learn about individual stimulus features.

Our behavioral analysis showed a main effect of target, which is consistent with Conjunctive learning and cannot be explained by Feature learning, Figure 2d. However, we observed an additional main effect of the number of features (single versus double). Subjects were faster, *F*(1,26) = 7.6, *p* = .01, 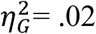, had a higher hit rate, *F*(1,26) = 27, *p* < .001, 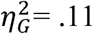, and made more false alarms, *F*(1,26) = 7.9, *p* = .008, 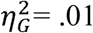, for two feature stimuli. Conjunctive learning cannot account for this effect, whereas Feature learning predicts this main effect, Figure 2d. This is because our models were initialized with zero values, which introduces a bias towards learning from target appearance relative to target non-appearance that disappears over time. This initialization improved the fit of all models. Because of this, the A feature in the Feature model has positive value despite being non-predictive of reward, leading to a higher value for conjunctions. Therefore, behavioral performance shows signatures of both Conjunctive and Feature learning.

Finally, we also observed an interaction between target association and the number of features, such that subjects showed an especially higher hit rate, *F*(1,26) = 9.7, *p* = .004, 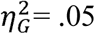, and higher false alarm rate, *F*(1,26) = 14.8, *p* < .001, 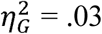, for AC-relative to B-trials. Only the Value Spread model, which mixes Feature and Conjunctive learning, can account for this interaction, Figure 2d. This interaction occurs in the Value Spread model because the bias towards learning about targets rapidly overwhelms the difference in AB+ and C+ due to feature learning, whereas the relatively slower initial learning about non-targets emphasizes the difference between AC- and B-. In sum, behavior shows patterns consistent with both Conjunctive and Feature learning, and the Value Spread model best accounts for the qualitative features of the data.

Response times exhibit base rate effects: if many recent trials required a response, the response on the subsequent trial is likely to be faster^28^. Such effects could be misinterpreted as feature learning if temporally adjacent target trials happen to share features. Our task design mitigated this concern because different trial types were randomly intermixed. Nonetheless, we formally tested whether such base rate effects could contribute to behavior and influence the performance of the Value Spread model. We augmented the Value Spread model with an additional base-rate learning mechanism. This agent learns the probability of a target, unconditional on the identity of the current stimulus. The values from this agent and the values from the Value Spread were entered as regressors on reaction time. The five parameters of this model (ω, *α*_*Value* *Spread*_, *β*_*Value* *Spread*_, *α*_*Base* *Rate*_, *β*_*Base* *Rate*_) were fit simultaneously. This addition did not significantly improve the likelihood of the Value Spread model, Wilcoxon *T* = 172, *p* = .14. Therefore, inclusion of base rate effects adds complexity without significantly improving the quality of model fits. Further, the regression betas on both the Value Spread RL value estimate and the Base Rate values were negative and significant (Value Spread RL *T* = 84, *p* = .002; Base Rate RL *T* = 136, *p* = .028), indicating that higher values from both agents are each associated with faster reaction times. Fits to the spread parameter, *ω* (mean: .41), indicate a relatively strong mixing of Conjunctive and Feature learning, even when accounting for response perseveration effects. We conclude that base rate effects did not qualitatively influence performance of the Value Spread model.

It has been proposed that subjects rely on hippocampal learning early in probabilistic reward tasks, and then transfer to using striatal learning over time^29^. We ran an additional control analysis to ensure that the performance of the Value Spread model across all trials did not reflect a transfer of learning from pure Conjunctive to pure Feature learning. We binned trials into early and late epochs, collapsing across runs. We examined the likelihood of the data for each epoch under the maximum likelihood parameter estimates for each model, Figure S1. We found an overall effect of epoch, *Z* = 4.7, *p* < .001, such that behavior was better fit by the models for later trials. However, we found no evidence of an interaction of model type (Feature versus Conjunctive) by epoch (early versus late), *p* > .2. These results indicate that performance of the Value Spread model likely does not reflect a transition from Feature to Conjunctive learning.

### 2.2 Striatal Prediction Error Analysis

Our behavioral analyses suggested subjects used a reinforcement learning strategy to acquire stimulus-outcome relationships, with learning best described by a model that spreads value updates among stimuli sharing features. Because striatal BOLD responses track reward PEs^30^, we predicted that these BOLD responses would co-vary with PEs derived from the Value Spread model. The key feature of this model is the spread of learning between stimuli sharing features, suggesting that subjects learn jointly about conjunctions and features. We sought to distinguish the contribution of Conjunctive learning, which learns independently about each stimulus, from that of Feature learning, which causes learning to spread across stimuli that share features. Such a distinction could emerge if the striatum integrated predictions arising from inputs from feature representations in sensory cortex and conjunctive representations in the hippocampus. A feature PE regressor was constructed from the Feature RL model. A conjunctive PE regressor was constructed by computing the difference between the feature regressor and PEs computed from the Conjunctive RL model. This regressor captures unique variance associated with PEs derived from a model that learns values for conjunctions (see Methods). In addition, constructing this regressor as a difference reduces the shared variation between the feature and conjunctive regressors. However, there remains shared variation between these regressors, *r*(118) *= -*.59, *p* < .001, and this shared variability reduces our ability to detect significant effects. Nonetheless, we found robust Feature PE responses in the bilateral medial caudate (whole-brain corrected threshold *p* < .05; Figure 2a). To confirm that this result was not driven by a higher response to targets than non-targets, we extracted single trial betas from an anatomical striatal mask^31^ crossed with a statistically-independent functional mask of Feature PE activation, and confirmed using a mixed-effects model with random-intercepts for subjects that both the target outcome, *t*(31) = 49.7; *p* < .001; *d*_*z*_ = 8.9, and the feature PE, *t*(31) = 5.7; *p* < .001; *d*_*z*_ = 1.02, contribute to the striatal outcome response. We use *d*_*z*_ to refer to Cohen’s d for one-sample tests. We next extracted parameter estimates from this ROI and observed these same voxels also showed evidence of a Conjunction PE response, *t*(31) = 4.1; *p* < .001; *d*_*z*_ = 0.72; Figure 2b. Note that this result is not driven by target coding because the Conjunction PE difference regressor is *anti*correlated with target occurrence (*r* = -.28), yet both show a positive relationship with striatal BOLD. Together, these results confirm that striatal BOLD tracked reinforcement learning PEs that mixed learning about conjunctions and features.

### 2.3 Pattern Similarity Analysis

We hypothesized the hippocampus formed conjunctive representations of task stimuli, which served as inputs to striatum for reinforcement learning. We used a pattern similarity analysis (PSA) to probe the representational content of hippocampus^32^. The PSA compares the similarity of activity patterns among different trials as a function of experimental variables of interest. We computed similarity matrices from the hippocampus, IFS, PRc, PHc and mOFC. The average similarity for each stimulus comparison is plotted in Figure S4.

We first tested whether representations of stimuli in the hippocampus remained stable across trials because to be useful for learning, a region must have consistent representations across presentations of a stimulus. We ran a regression analysis on PSA matrices to assess the similarity among representations from different presentations of a stimulus. We tested the significance of each effect by permuting the PSA matrices 10,000 times to build a null distribution of regression coefficients. All ROIs had significantly higher similarity for repetitions of the same stimulus (within-stimulus similarity) than for pairs of different stimuli (between-stimulus similarity; all *p* < .001, FDR corrected) except for the mOFC, *p* > .3. Therefore, all ROIs except for the mOFC have representations that are driven by the stimulus. Across-region comparisons showed the hippocampus had stronger within-stimulus coding than PRc, *p* < .001, PHc, *p* < .001, IFS, *p* < .001 and mOFC, *p* < .001, FDR corrected, Figure 3a, indicating the hippocampus had the most stable representations of task stimuli.

**Figure 3.**
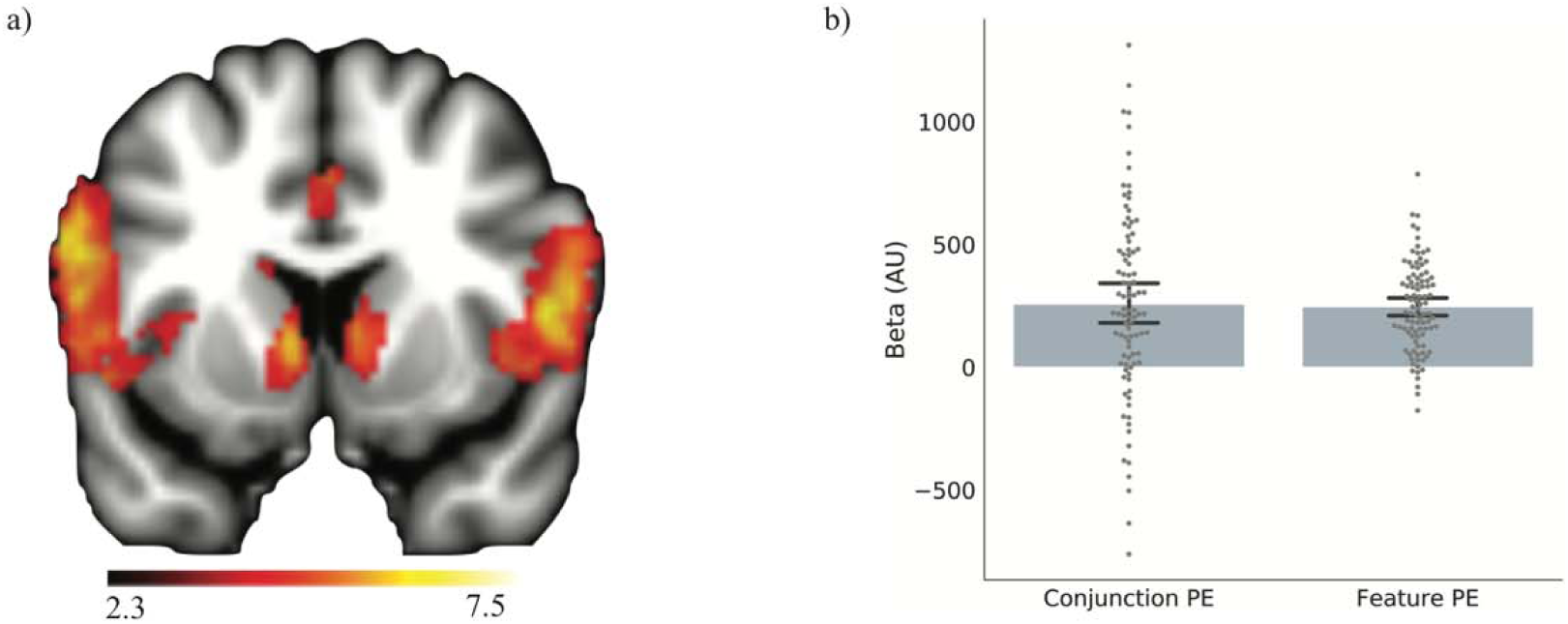
Striatal error response. a) Regions responsive to PEs from Feature RL model (whole-brain analysis; *p* < .05). b) An ROI analysis of striatum showed that voxels with responses that scaled with PEs from the Feature RL model also scaled with PEs from the Conjunctive RL model. The Feature PE bar is a statistically independent depiction of the striatal response in (a). The Conjunction PE bar shows that errors from a conjunctive learning system correlated with striatal BOLD above and beyond errors from a feature learning system. Dots correspond to individual runs with the subject intercepts removed.

Our central hypothesis was that the hippocampus, not PRc, PHc, mOFC nor IFS, would form conjunctive representations of stimuli. Representations of stimuli that shared features (AB and B) should be pattern separated, and therefore less correlated with one another, in hippocampus but should be more correlated with one another in cortical regions like PRc and PHc that provide inputs to the hippocampus. Our task intentionally included speeded responses and probabilistic outcomes, features that are known to bias subjects away from employing rule-based strategies^33^. As a result, predicted that the IFS should not have pattern-separated representations of conjunctions. Finally, because mOFC is not associated with pattern separation, we predicted that the mOFC would not have pattern separated responses. We tested whether the representational structure in each ROI was more similar for stimuli sharing common features than for stimuli that lacked feature overlap [(AB, AC), (AB, B), (AC, C)] versus [(AB, C), (AC, B), (B, C)]. We note that this analysis is orthogonal to the previous within-stimulus analysis and provides an independent test of stimulus coding fidelity. We also note that all our stimuli, including single-feature stimuli, are in reality conjunctions of features because they are experienced in our task context (i.e., for common task context X; stimuli are truly ABX, ACX, BX, CX). Therefore, rather than testing for differences between conjunctive and non-conjunctive stimuli, our overlap regressor tests for similarity in representations between conjunctive stimuli that share a salient feature versus those that do not. All control ROIs showed a significant effect of overlap, PRc: *p =* .01, PHc: *p* = .009, IFS: *p* < .001, and mOFC: *p* < .001, FDR corrected, Figure 3b, but the hippocampus did not, p > .3. Critically, the hippocampus showed significantly lower representational overlap than PRc, *p* = .015, PHc, *p* = .015, IFS, *p* = .002 and mOFC, *p* = .002, all FDR corrected. Control analyses ruled out potential confounds arising from feature hemifield and reproduced these findings using a parametric mixed-effects model (Supplemental Information). There is a correlation between the overlap regressor and the effect of response, i.e., comparisons between stimuli with the same target outcome versus different target outcome (*r* = .09). We included nuisance regressors to control for this effect. In addition, this correlation can only introduce a spurious increase in our overlap measure if cortical ROIs are *more* similar when the outcomes are *different* (because there are more pairs with different outcomes in the non-overlap condition; [(AB+, AC-), (AB+, B-), (AC-, C+)] versus [(AB+, C+), (AC-, B-), (B-, C+). Empirically, the effect of response is positive in all regions except mOFC (Figure S5), indicating residual effect of response coding not accounted for by our regression approach would lead to an underestimate in our main finding of interest. In sum, relative to the control ROIs, the hippocampus formed more pattern-separated conjunctive representations of stimuli.

Hippocampal representations of conjunctions could serve as inputs to the striatal reinforcement learning system. If this is the case, then variability in the formation of pattern-separated conjunctive representations in hippocampus should correlate with striatal learning about conjunctions. When the hippocampus demonstrates relatively more pattern-separated representations, the striatal error signal should more strongly track PEs estimated from the Conjunctive RL model. To examine this relationship, we fit a mixed effects model of the conjunctive component of the striatal PE, with subject as a random intercept and hippocampal overlap as a random slope. The hippocampal overlap term was *negatively* related to the striatal conjunctive PE, *t*(31) = −3.43, *p* = .003, *d*_*z*_ = −0.62, Figure S2. The conjunctive PE represents variance explained over-and-above the effect of feature PE and is therefore a more sensitive measure of the degree of conjunctive PE coding in the hippocampus. As expected, there was no relationship between hippocampal overlap and the striatal feature PE, *p* > .2, and this finding suggests that general signal quality fluctuations did not contribute to the effect. Control analyses showed that this result persisted even when accounting for inter- and intra-individual differences in how well subjects learned (Supplemental Analyses), suggesting that it was not entirely driven by how much attention subjects paid to the task. However, attention is likely to be an important driver of both hippocampal pattern separation and striatal learning. Given our model that representations in both sensory cortex and HPC project to the striatum to influence learning, a relative increase in overlapping representations in any of these regions should be associated with a reduced conjunctive component of the striatal prediction error. We observed similar relationships in our medial temporal lobe (MTL) cortical ROIs and IFS, but not OFC (Supplemental Information). Finally, we observed a *positive* relationship between the strength of within-stimulus similarity in the hippocampus and striatal conjunctive PE, *t*(31) = 2.49, *p* = .013, *d*_*z*_ = 0.45, although this result depended on the exclusion of an outlier subject (Figures S2). Again, this finding suggests that the results were not driven by general signal quality issues, as stimulus identity and stimulus overlap showed opposing relationships to the conjunctive PE in the predicted direction. In sum, the more the hippocampus established pattern-separated representations of stimuli, the more striatal error signals reflected learning signals arising from a conjunctive state space.

We next tested whether there was a relationship between hippocampal overlap and behavior. We computed an index that measures the extent to which subjects used Conjunctive learning (see Methods) for each run of subjects’ behavior. We fit a mixed-effects model of this measure with random intercepts for subjects and included a nuisance covariate that measured how well subjects learned in each run, relative to chance. This helps to ensure that the relationship is not due to fluctuations in participant engagement. Contrary to our predictions, we did not find any relationship between overlap in any of our ROIs and this measure. Variability between runs in this measure may be too large to detect fine scale relationships with behavior. We next examined whether conjunctive learning was related to univariate signal magnitude, and found that runs with stronger hippocampal activity were also runs with the most conjunctive learning, *t*(31) = 3.1, *p* = .01, *d*_*z*_ = 0.56, FDR corrected. This relationship was nonsignificant in other ROIs, all *p* > 1. Therefore, univariate signal in the hippocampus is predictive of conjunctive learning.

### 2.4 Representation of stimuli according to their association with the target

How the hippocampus represents stimuli with similar associations is a question of current debate. Computational models suggest that the hippocampus supports feedback learning by representing stimuli with similar outcomes more similarly^34^. This grouping facilitates responding, while also allowing generalization of knowledge across related items^27,35,36^. However, an alternative perspective maintains that in order to maintain distinct representations of related items, the hippocampus orthogonalizes stimuli with related outcomes *more* strongly. Recent work has shown that learning drives representations of stimuli with similar outcomes apart in the hippocampus, resulting in representations of similar items that are even more distinct than representations of unrelated items^37,38^. We conducted an exploratory analysis to assess how our ROIs represented stimuli according to the strength of their association with the target, Figure 5a. We used the values from the Value Spread model to compute a value regressor that was higher when stimulus-target associations are more similar. This regressor had a negative relationship to hippocampal pattern similarities, which shows that the hippocampus represents stimuli with similar values more distinctly, *p* < .001, FDR corrected. We also observed this relationship in the PRc, *p* < .001 and PHc, *p =* .001. In contrast, the mOFC showed a positive effect of value, representing stimuli with similar association to the target more similarly, *p* = .001, FDR corrected. Similar to the overlap analysis, we also found a marginal effect of higher striatal conjunctive PE on runs in which trials with similar outcomes are represented more dissimilarly in the hippocampus, *t*(31) = −2.24, *d*_*z*_ = −0.40, *p* = .063, and PRc, *t*(31) = −2.48, *p* = .063, *d*_*z*_ = −0.45, FDR corrected.

**Figure 4.**
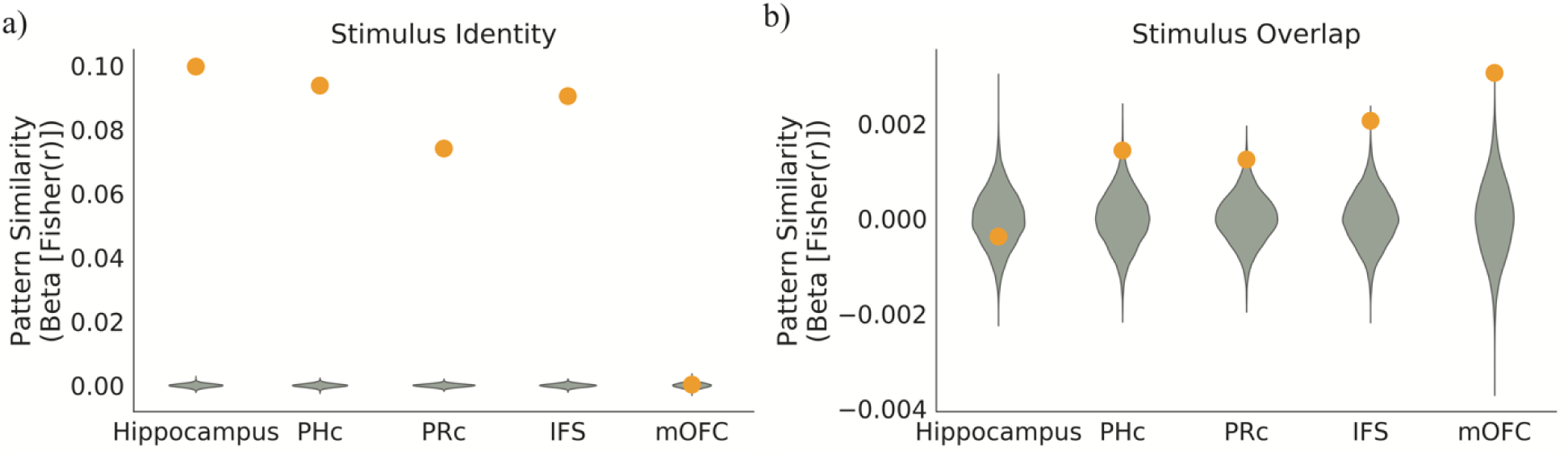
Pattern similarity analysis. a) A regression analysis on PSA matrices showed strong within-stimulus coding in all ROIs except mOFC, and within-stimulus coding was significantly stronger in hippocampus relative to other regions. The y-axis shows regression weights from a within-stimulus regressor on the PSA matrix of each ROI. b) PHc, PRc, IFS and mOFC showed increased similarity for pairs of stimuli that shared features and significantly more similarity for these pairs than the hippocampus, consistent with pattern-separated representations in the hippocampus. Green violins show the null distributions of regression coefficients from 10,000 randomly permuted PSA matrices. The y-axis shows regression weights from an overlapping-versus-non-overlapping stimuli regressor on the between-stimuli correlations from the PSA matrix of each ROI.

**Figure 5.**
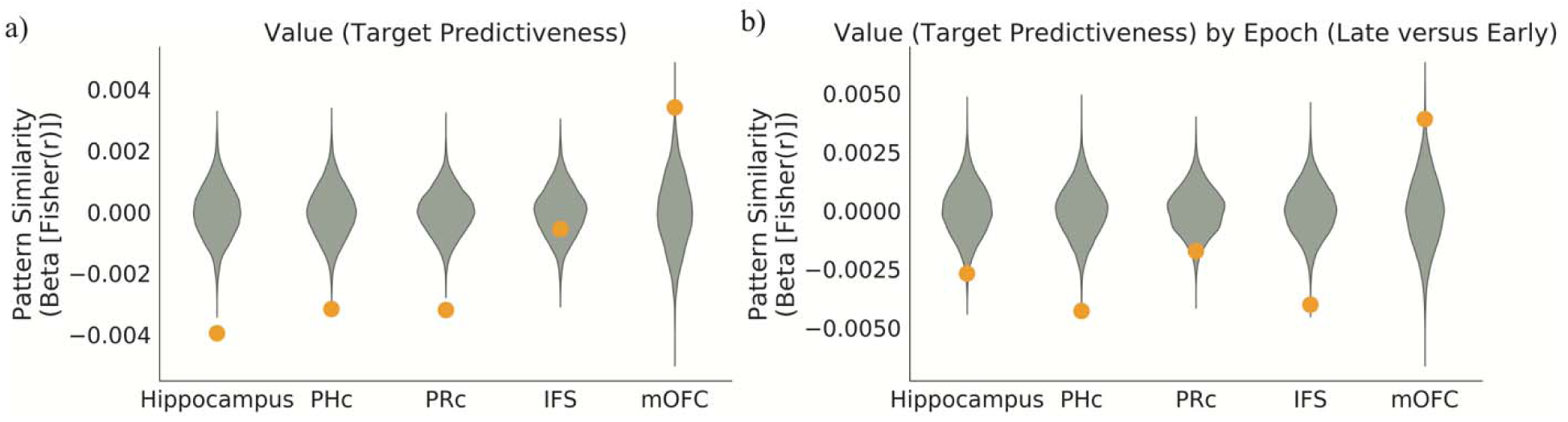
Pattern similarity analysis of stimulus value. a) A regression analysis on PSA matrices showed strong that stimuli with similar associations to the target are further apart in representational space in the hippocampus, PHc, and PRc. In contrast, they are closer together in representational space in the mOFC. The y-axis shows regression weights from value similarity regressor on the PSA matrix of each ROI. b) Hippocampus, PHc, IFS and PRc representations for stimuli with similar associations with the target moved further apart in representational space over the course of a run. In contrast, in mOFC, representations became more similar. Green violins show the null distributions of regression coefficients from 10,000 randomly permuted PSA matrices. The y-axis shows regression weights from the interaction of the value similarity regressor with a regressor that encodes comparisons between trials late in learning versus comparisons between trials early in learning.

Because previous work showed that representations became progressively more dissimilar over the course of learning, we tested whether this effect interacted with task epoch. For this analysis, we constructed a new model with an additional interaction term between stimulus value and an epoch regressor, which was positive for comparisons between stimuli late in the run and negative for comparisons between stimuli early in the run. We found an interaction in the hippocampus, *p* =.01, FDR corrected, such that the representational distance between stimuli with similar values increased over the run, Figure 5b. We also observed this effect in PHc, *p* < .001, IFS, *p* < .001 and PRc, *p* = .035, FDR corrected. In contrast, in mOFC, we found the opposite: representational distance for stimuli with similar values decreased over the run, *p* = .01, FDR corrected.

### 2.5 Representational Content Analysis

The previous analyses show that the hippocampus has the most distinct representations of stimuli that share features among our regions of interest. However, the demonstration of no significant increase in similarity of hippocampal representations for feature-sharing stimuli begs for a more direct test of pattern separation. To directly test this hypothesis, we probed the content of hippocampal and cortical ROI representations using estimates of categorical feature coding acquired from independent localizer data. If hippocampal conjunctive representations are pattern separated from their constituent features, then they are not composed of mixtures of representations of those features^39,40^ (Figure 4a). Unlike high-level sensory cortex, the hippocampal representation of {face and house} would not be a mixture of the representation of {face} and {house}^39^. We predicted that the hippocampal representations of two-feature stimuli ({face and house} trials) in our learning task should be dissimilar from representations of faces and houses in the localizer. Hippocampal representations of one-feature trials ({face} trials), which are less conjunctive because they contain only one task-relevant feature, should be more similar to representations of the same one-feature category (e.g., faces) in the localizer. In contrast, cortical representations of both two-feature and single-feature trials should be similar to representations of their corresponding features in the localizer (Figure 4a). We predicted that hippocampal representations would be less similar to feature templates than cortical ROIs, and that only the hippocampus would show would less similarity for two-feature than single-feature trials.

We correlated the patterns in each ROI with the corresponding localizer feature templates (Methods) but were unable to detect reliable feature responses from IFS or mOFC in the localizer data^41^. We found significant similarity among task patterns and feature templates for all conditions, *p* < .001, FDR corrected, except for hippocampal responses to conjunctive stimuli, *p* = .116, Figure 6. The hippocampus had lower similarity to feature templates than PRc, *p* < .001, and PHc, *p* < .001. This effect was not likely driven by regional signal quality differences, as the hippocampus had the strongest within-stimulus coding (Figure 4a). The hippocampal feature template was more similar to the response to a single-feature stimulus than to a two-feature stimulus, *p* < .001, consistent with a gradient in pattern separation as the number of task-relevant features increased. This effect was larger in the hippocampus than either PRc, *p* < .001, or PHc, *p* < .001. We confirmed these results using a parametric mixed effects analysis and also performed a control analysis to verify the results were not driven by stimulus-general activation (Figure S6). Unexpectedly, we observed that similarity in PRc and PHc was stronger for two-feature than for single-feature stimuli, both *p* < .001; in the mixed effects model, this result was marginal in PRc and nonsignificant in PHc and should be interpreted with caution.

**Figure 6.**
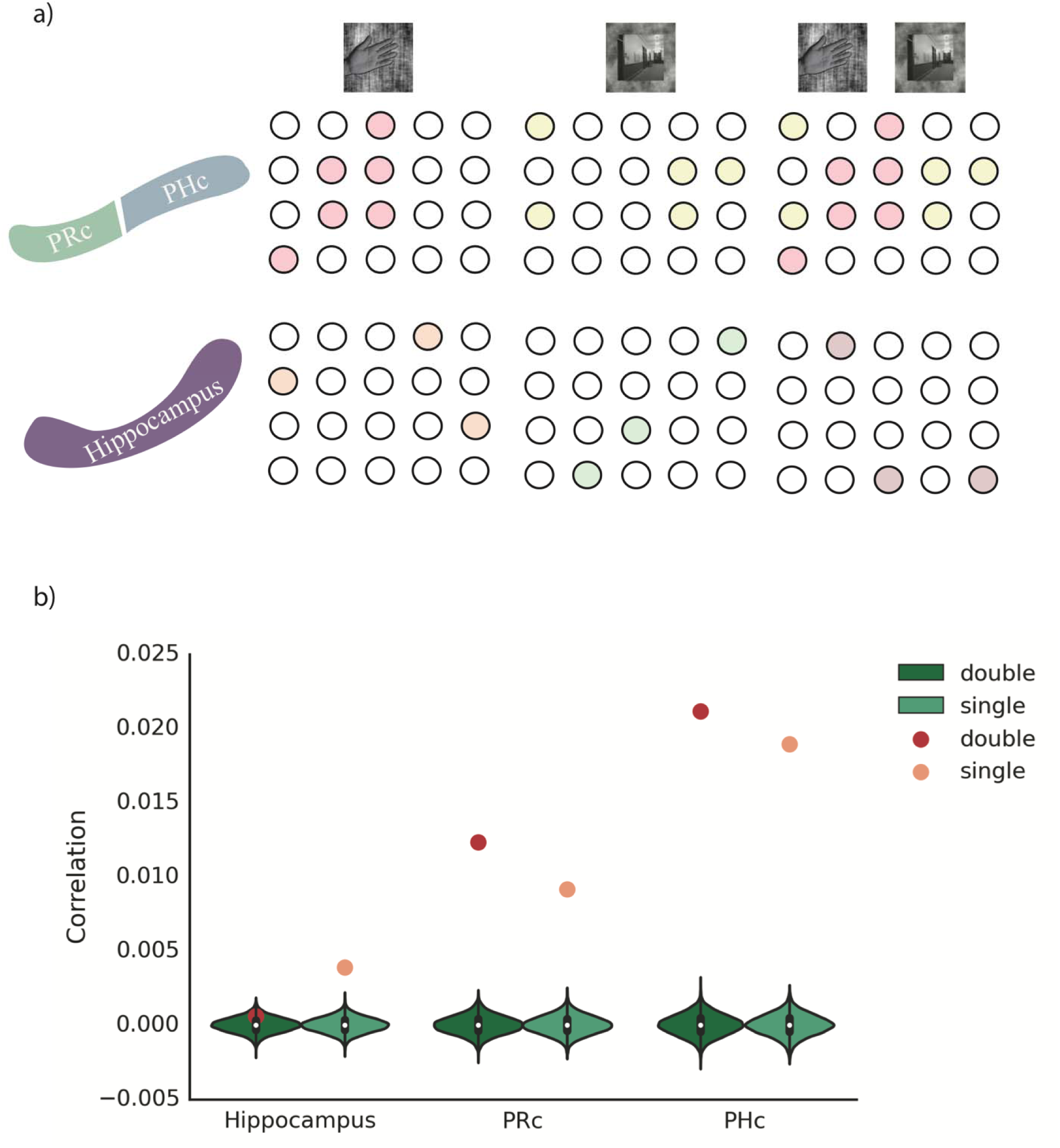
Representational content analysis. a) Neural predictions: top panel is putative neural ensembles in high-level sensory cortex (parahippocampal, PHc; perirhinal, PRc) for task stimuli^42^. Two-feature stimulus should be represented as union of responses to component features. The lower panel shows putative neural ensembles in the hippocampus; the neural representation of two-feature conjunctions should be orthogonal to responses to its component features. b) Hippocampal representations were less similar to feature templates than PRc and PHc representations, consistent with increased conjunctive coding. In the hippocampus, representational similarity to templates was higher for single-feature than two-feature stimuli, consistent with increased pattern separation for stimuli with multiple task-relevant features. PRc and PHc showed increased similarity for two-feature relative to one-feature stimuli.

## 3 DISCUSSION

We tested whether pattern separation in hippocampus enabled learning stimulus-outcome relationships over multifeatural stimuli. We used a novel reinforcement learning task that required learning over non-spatial conjunctions of features. The hippocampus encoded stable representations across repetitions of a stimulus, and conjunctive representations were distinct from the representations of composite features. The hippocampus showed stronger evidence for pattern-separated conjunctive representations than PRc, PHc, IFS and mOFC. Hippocampal coding was also related to PE coding in the striatum. Our results suggest that the hippocampus provides a pattern-separated state space that supports the learning of outcomes associated with conjunctive codes.

Our finding of overlapping representations of conjunctions that share features in PRc and PHc, in combination with our finding of mixed conjunctive and feature learning in both behavior and the striatal error response, suggests that feature representations influence learning even when they do not benefit performance. The striatum receives inputs from diverse cortical and subcortical areas and may integrate predictions from systems that represent the environment in different ways (e.g., conjunctive versus feature)^43,44^. It seems reasonable that, over longer training periods, subjects would learn to down-weight the influence of cortico-striatal synapses representing uninformative features. Attention may support this process by influencing the relative strength of representations of conjunctions versus features^24^. Future work should examine learning over longer timescales in order to evaluate how attention may influence cortical and hippocampal processing in order to prioritize learning about more informative aspects of the environment.

Lesion studies showing dissociations between the hippocampus and striatum in learning^45^ along with some imaging studies demonstrating a negative relationship between hippocampal and striatal learning signals^29^ have led to the hypothesis that these regions compete during learning and that learning transfers from hippocampal to striatal systems over time. By contrast, other evidence document cooperative interactions. For example, neurons in the striatum represent spatial information derived from hippocampal inputs^46^ and contextual information in the hippocampus drives the formation of conditioned place preferences via its connection to ventral striatum^43,47^. These findings, in concert with our own, support a model in which hippocampal information about spatial contexts, location, or conjunctions serve as inputs for striatal associative learning.

Our finding that the hippocampus represents stimuli with similar outcomes more differently, and that this representational distance increases during learning, is consonant with recent demonstrations of repulsion of hippocampal representations of related memories in navigation^38^ and in relational learning^37^. These findings, together with the present results, suggest that the hippocampus dynamically increases the representational distance between overlapping experiences that may otherwise be subject to integration in cortical circuits. However, our results are inconsistent with a related study investigating hippocampal involvement in value-based decision-making^48^. Barron et. al pre-exposed subjects to individual foods (e.g, tea and jelly) and then asked them to simulate the value of a food that combines the individual foods (e.g, tea flavored jelly). They observed repetition suppression for compounds (AB) that were preceded by their components (A or B), indicating a representation of the features as part of the compound. The inconsistency between our results could arise because the task in Barron et. al required subjects to integrate singletons to imagine the value of a conjunction, whereas our task required subjects to distinguish between singletons and conjunctions. This distinction is reflected in a large literature showing that the hippocampus maintains distinct representations of related experiences^49^ while also maintaining a representation of the relationships between them^35,36,50^. Goal-directed attention may play an important role in determining which aspect of hippocampal function comes to dominate the hippocampal representation^51^. In our data, attention could be a mediating variable that drives both the hippocampal response, striatal learning signals and fluctuations in behavioral performance. Our finding of a selective relationship between hippocampal BOLD and behavioral performance against a simple model in which wholesale fluctuations in participant engagement underlie our effects. Rather, attention may selectively influence the fidelity of hippocampal representations of conjunctions in order to support conjunctive learning.

Our results complement and extend a recent investigation of the role of the hippocampus in conjunctive learning^52^. In this task, subjects were required to learn the relationship between conjunctions of cues and a weather outcome. They observed a univariate relationship between the hippocampus BOLD and the degree of conjunctive learning, as well as a correlation between hippocampus and nucleus accumbens that relates to conjunctive learning. In addition to these univariate effects, they also observed a within-stimulus similarity effect in the hippocampus but did not investigate stimulus overlap nor the representations of other cortical ROIs. Our results therefore provide a unique demonstration that the hippocampal code is more pattern separated than other cortical ROIs during conjunctive learning.

We contend the hippocampus forms representations of conjunctions of features that are reinforced via dopamine release on hippocampal-striatal synapses, but the hippocampus could form a representation of the temporal sequence of task events^34^. In AB+ trials, AB could trigger a representation of the target in the hippocampus, and this target representation could then feed into the striatum or prefrontal cortex to drive responses. This model is similar to the idea that the hippocampus encodes a “successor representation” for reinforcement learning^53^ in which the target representation occurs in proportion to the probability of each stimulus preceding the target. The hippocampus-to-striatum connectivity and successor representation explanations of our results differ in mechanism, but share the requirement of a conjunctive representation in the hippocampus. Future work should directly test the role of hippocampal sequence representation in reinforcement learning.

We did not find evidence for the coding of stimulus identity in the mOFC; however, we found that the mOFC represented stimuli with similar strength associations to the target more similarly. Further, this effect increased from early to late in runs, such that mOFC representations of stimuli with similar target associations moved closer together in representational space. These findings support a model in which the mOFC representation is primarily driven by outcome prediction and outcome representation (Figure S5). This finding is inconsistent with the strongest version of a recent theory suggested that the OFC supports a state space representation in learning tasks^21^. However, that theory suggests that the OFC is particularly important for representing aspects of the state space that are unavailable in the immediate sensorium, such as information held in working memory. Our task has no requirement for such a function. It is also possible that OFC state representations are structured so as to be most useful for behavior. Specifically, the OFC could represent just two states corresponding to different action policies ({AB+, C+} versus {AC-, B-}). We prefer the interpretation that the mOFC representations in our task are driven by the subjective value of the stimuli, because the target is associated with the possibility for reward^18^. Finally, our distinct findings between mOFC and hippocampal representations echo recent investigations of context-based decision in rodents showing that the hippocampal representation is primarily driven by the context^54^, whereas the orbitofrontal cortex representation is primarily driven by reward value^55^.

The circuit properties of the hippocampus allow it to rapidly bind distributed cortical representations of features into orthogonalized conjunctive representations. Hippocampal pattern completion, triggered by partial cues, along with recurrent outputs back to sensory cortex allow the hippocampus to reactivate the ensemble of event features that constitute the retrieval of an episodic memory^2,56^. Dense inputs to the striatum suggest hippocampal representations could also form the basis for associative learning over conjunctive codes. Our results extend the role of the hippocampus to include building conjunctive representations that are useful for striatal outcome and value learning.

## 4 METHODS

### 4.1 Data and Software Availability

All MRI and behavioral data will be made available at OpenfMRI, and all analysis code will be made available on GitHub prior to publication. Key resources are listed in Table S2.

### 4.2 Experimental Model and Subject Details

The study design and methods were approved by the Stanford Institutional Review Board. Forty subjects provided written informed consent. Data from eight subjects were excluded from analyses: One ended the scan early due to claustrophobia; three had scanner-related issues that prevented reconstruction or transfer of their data; two had repeated extreme (>2mm) head movements across most runs; and two subjects demonstrated extremely poor performance, as indexed by less than $2.50 of earnings (see below for payment details). Note that our task is calibrated to the individual subjects’ practice data in such a way that a simple target detection strategy would be expected to earn $7.50, and any effort to learn the task should improve on these earnings. This left 32 subjects in the analysis cohort, 19 females, mean age 22.1 years, SD 3.14, range 18 to 29. Due to an error, behavioral data for one subject were lost; thus, while her imaging data were included in fMRI analyses, all behavioral analyses were conducted with a sample of 31 subjects.

### 4.3 Task

Subjects performed a target detection task in which performance could be improved by learning predictive relationships between visually presented stimuli and the target. The target appeared 70% of the time for two-feature stimulus AB as well as the single-feature stimulus C, and 30% of the time for two-feature stimulus AC as well as the single-feature stimulus B. The task bears strong similarities to the “ambiguous feature discrimination task” used to study rodent learning^23^. Subjects were instructed that they would earn 25¢ for each correct response, lose 25¢ for each incorrect response, or no money for responses that are slower than threshold or missed.

Response time (RT) thresholds were calibrated for each subject so that responses initiated by perception of target onset would lead to success on 50% of trials. This procedure incentivizes subjects to learn predictive information in order to respond more quickly. Before scanning, subjects performed a simplified target-detection trial in which they responded to a probabilistic target with no predictive relationships between the cue and target. During this session, RT thresholds were adjusted by 30 ms increments on each trial (fast-enough responses reduced the threshold while too-slow responses increased it). During the task, we continued to make smaller changes (10 ms) so that the threshold could change if subjects became progressively faster. Earning rewards on more than 50% of trials required anticipating target onset based on the preceding stimulus (A, B, AB, or AC). Subjects were instructed that in order to earn the most money, they should learn which stimuli predicted the target and respond as quickly as possible, even if the target has not yet appeared. These instruction were meant to bias subjects towards an instrumental learning strategy, rather than an explicit rule-based learning strategy^33^.

Subjects performed one practice run and were instructed on the types of relationships they might observe. During fMRI scanning, subjects performed three runs of the task. Each run consisted of 10 trials for each stimulus (AB+, AC-, B-, C+), resulting in 40 total trials per run. Features A, B, and C were mapped to a specific house, face, and body part image for the duration of the run. Subjects were not pre-exposed to the specific stimuli. The category-to-stimulus mapping was counterbalanced across runs, resulting in each visual category being associated with each feature type (A, B or C) over the course of three runs. The counterbalancing of category-to-stimulus mapping ensures that any carry-over effects of learning across runs can only have a detrimental and noisy effect on learning that would work against our hypotheses. In addition, we verbally emphasized that mappings changed between runs. Further, different participants saw different stimuli within each category and different stimuli across runs. Each subject encountered the same pseudo-random trial sequence of both stimuli and targets, which facilitated group modeling of parametric prediction error effects. Features could appear on either the left or the right of a fixation cross, with the assignment varying randomly on each trial. For single-feature stimuli, the contralateral location was filled with a phase-scrambled image designed to match the house/face/body part features on low-level visual properties. The target stimulus was a car image and was consistent across all trial types. On trials where the target appeared, it did so 600 ms after the onset of the visual cues. Inter-trial intervals and the interval between the stimulus/stimulus+target and feedback were taken from a Poisson distribution with a mean of 5 s, truncated to have a minimum of 2 s and maximum of 12 s. Visual localizer task details are described in *SI Methods*.

### 4.4 Behavioral Analysis

We used reaction time data from subjects to infer learning in the task, an approach that has been used successfully in a serial reaction time task^57^. Log-transformed reaction times were fit with linear regression, with the difference with the difference that we jointly fit the parameters of the regression model and the parameters of a reinforcement learning model. Specifically, we modeled reaction time with a value regressor taken from a reinforcement learning model:

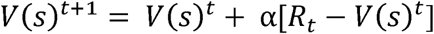

where *R*_*t*_ is an indicator on whether or not the target appeared, α is the learning rate, and *s* is the state. These values represent the strength of association between a stimulus and a target/outcome, and are not updated based on the reward feedback, which also depends on whether the subject responded quickly enough. The values we measured are more relevant for learning because they correspond to the probability that the subject should respond to the stimulus. We constructed values from three different models that made different assumptions about the underlying task representation. For the Conjunctive and Value Spread models, the state corresponded to the current stimulus *s* 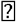{B, C, AB, AC}. For the Feature model, the states were single features *s* 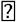{A, B, C} and in two-feature trials (e.g., AB), the value was computed as the sum of the.individual feature values (e.g., V(AB) = V(A) + V(B)), and both feature values were updated after feedback. Finally, the base rate learning maintained a single state representation for all four stimuli.

The Value Spread RL model was a variant of the Conjunctive model in which a portion of the value update blends onto the overlapping stimulus:

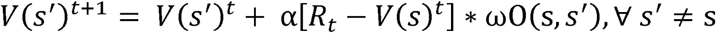

where O(*s,s’*)is an indicator function that is equal to 1 when the two stimuli share a feature and 0 otherwise, and ω is a spread parameter that controls the magnitude of the spread. For example, if the current stimulus is AB, a proportion of the value update for AB would spread to B and to AC. We allowed value to spread between any stimuli sharing features (e.g., an AB trial would lead to updates of both AC and B). This approach reflects the fact that not only will conjunctions activate feature representations in cortex, but features can activate conjunctive representations, a property that has been extensively studied in transitive inference tasks^58^. We fit the Conjunctive RL model, the Feature RL model, and the Value Spread RL model using Scipy’s minimize function. The linear regression weights were fit together with the parameters of the learning model. We calculated likelihoods from the regression fits using the standard regression formula that assumes normally distributed errors (*σ*^2^):

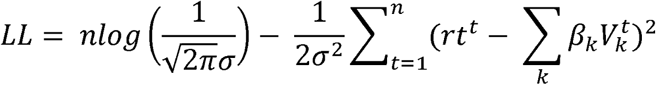

Where *n* is the number of trials, the 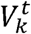 are *k* different estimates of value at trial t, and the *β*_*k*_ are regression coefficients. Finally, we fit a null No Learning model with no value regressor (and therefore fits to only mean RT). Model comparison procedures are described in detail in

#### Supplemental Methods

In order to analyze the relationship between striatal BOLD and model-derived prediction error, we modeled the Feature model prediction error as well as the difference between the Conjunctive model prediction error and the Feature model prediction error. This subtraction approach has three advantages: 1) it reduces the shared variance considerably from modeling the two prediction errors (feature and conjunction, r(118) = .79, Note that the correlation flips sign because of the subtraction); 2) it provides a stronger test about whether conjunctive representations contribute to striatal error responses, because that contribution must be over and above the contribution of a feature learning model; and 3) together, the two regressors combined are a first-order Taylor approximation to a hybrid model that weighs contributions of a conjunctive and a feature learning mechanism. This hybrid model is conceptually very similar to the Value Spread model, although distinct in minor aspects. This means that we can model the data by supposing a hybrid mechanism, as we found in behavior, while also separately examining components of that hybrid mechanism.

In order to compute our conjunctive learning index we computed the likelihood of the data for each run and subject under the maximum likelihood parameters for the Value Spread model and the Conjunctive model. We computed the difference in likelihood between the models. This measure reflects the extent to which the Conjunctive model accounts for the data better than the Value Spread model. For our nuisance regressor, we computed the difference in likelihoods between the Value Spread and the no learning model.

## 4.5 fMRI Modeling

fMRI acquisition and preprocessing as well as ROI selection procedures are described in detail in SI Methods and ROIs are depicted in Figure S3. Analysis was conducted using an event-related model. Separate experimental effects were modeled as finite impulse responses (convolved with a standard hemodynamic response function). We created separate GLMs for whole brain and for pattern similarity analyses (PSA). For the whole brain analysis, we modeled the 1) stimulus period as an epoch with a duration of 2 s (which encompasses the stimulus, target and response) and 2) the feedback period as a stick function. In addition, we included a parametric regressor of trial-specific prediction errors extracted from a Feature reinforcement learning model that learns about A, B, and C features independently, without any capacity for learning conjunctions. In addition, we included a regressor that was computed by taking the difference between the Feature RL errors and the Conjunctive RL model errors. This regressor captured variance that was better explained by Conjunctive RL prediction errors than by Feature RL prediction errors. Both parametric regressors were z-scored and were identical across all subjects. We included nuisance regressors for each slice artifact and the first principal component of the deep white matter, which captures residual nuisance components of the whole-brain signal. Following standard procedure when using ICA denoising, we did not include motion regressors as our ICA approach is designed to remove motion-related sources of noise. GLMs constructed for PSA had two important differences. First, we did not include parametric prediction error regressors. Second, we created separate stimulus regressors for each of the 40 trials. Models were fit separately for individual runs and volumes of parameter estimates were shifted into the space of the first run. Fixed effects analyses were run in the native space of each subject. We estimated a nonlinear transformation from each subject’s T1 anatomical image to the MNI template using ANTs. We then concatenated the functional-to-structural and structural-to-MNI transformations to map the fixed effects parameter estimates into MNI space. Group analyses were run using FSL’s FLAME tool for mixed effects.

## 4.6 PSA Analysis

We were interested in distinguishing the effect of different experimental factors on the representational similarity matrices (PSM). PSM preprocessing is described in SI Methods. We constructed linear models of each subject’s PSM. We included main regressors of interest as well as several important regressors that controlled for similarities arising from task structure. The main regressors of interest were 1) a “within-stimulus similarity” regressor that was 1 for pairs of stimuli that were identical and 0 otherwise and 2) an “overlap” regressor that was coded as 1 for pairs of stimuli that shared features, −1 for those that did not, and 0 for pairs of the same stimuli. We also included exploratory regressors of interest 3) a “prediction error” regressor that was computed as the absolute value of the difference in trial-specific prediction errors, extracted from the Value Spread reinforcement learning model, between the two stimuli (Figure S5), 4) a “value” regressor that was computed as the absolute value of the difference in trial-specific updated values, extracted from the Value Spread reinforcement learning model, between the two stimuli (Figure 5). We included two nuisance regressors that model task-related sources of similarity that are not of interest: 5) a “response” regressor that was coded as 1 for stimuli that shared a response (both target or both non-target) and −1 otherwise, 6) a “target” regressor that was coded as 1 for stimuli that both had a target, −1 for both non-target, and 0 otherwise. Finally, we included nuisance regressors for 7) the mean of the runs and 8) two “time” regressors that accounted for the linear and quadratic effect of time elapsed between the pair of stimuli and 9) the interaction between the time regressor and the within-stimulus similarity regressor. We included this last interaction because within-stimulus similarity effects were by far the most prominent feature of the PSA, and temporal effects were therefore more likely to have larger effects on this portion of the PSMs. We included prediction error (3) and value (4) regressors because we were interested in exploratory analyses of these effects based on theoretical work suggesting that the hippocampus pattern separates stimuli based on the outcomes they predict34. Both the value and the prediction error regressors were orthogonalized against the response and target regressors, thereby assigning shared variance to the regressors modeling outcomes. All regressors were z-scored so that their beta weights could be meaningfully compared. Correlations, our dependent variable, were Fisher transformed so that they followed a normal distribution. To assess the significance of the regression weights as well as differences between regions, we compared empirical regression weights (or differences between them) to a null distribution of regression weights (or differences between them) generated by shuffling the values of the PSA matrices 10,000 times. In addition, we fit a linear mixed effects model using R with subject as a random intercept, ROI as a dummy code with hippocampus as the reference, and ROI by task interactions for each of the above regressors. Using random slopes resulted in convergence errors, and so we did not include them. By using both a parametric and a nonparametric approach to assessing our data, we gained confidence that our results are robust to differences in power between different statistical analysis techniques due to outliers or violations of distribution assumptions.

## 6.7 Representational Content Analysis

Data from the localizer task were preprocessed and analyzed in the same manner as the main task data. GLMs were constructed for each run and included a boxcar regressor for every miniblock with 4-s width, as well as a nuisance regressor for the targets, each slice artifact and the first principal component of the deep white matter. To compute template images, we computed the mean across repetitions of each stimulus class (face, place, character, object, body part). For the representational content analysis depicted in Figure 4, we computed the correlations as follows: Assume that A is a face, B is a house, and C is a body part. For “single-feature” stimuli, we computed the similarity of B trials with the house template and C trials with the body part template. For “two-feature” stimuli, we computed the similarity of AB trials with the house template and AC trials with the body part template. Therefore, within each run, the task-template correlations for AB and B (and AC and C) were computed with respect to the same feature template. This means that any differences between AB and B (or AC and C) correlations reflect differences in the task representations, rather than potential differences in the localizer representations. We repeated this for each run’s stimulus category mappings.

## 5 AUTHOR CONTRIBUTIONS

Conceptualization, ICB and SMM; Methodology, ICB; Software, ICB; Formal Analysis, ICB; Data Curation, ICB; Writing Original Draft, ICB; Writing, Review & Editing, SMM and ADW.; Funding Acquisition, ICB and SMM; Supervision, SMM and ADW.

## 6 ACKNOWLEDGEMENTS

We thank the NSF GRFP (ICB), NSF IGERT (ICB) and Stanford Innovation Grants (ICB). We also thank Kim D’Ardenne for significant editing and Stephanie Gagnon, Karen LaRocque, Alex Gonzales, Anna Khazenzon and Yuan Chang Leong for their feedback.

